# Proteomic adaptations in the kidney reveal orchestration of local and secreted antimicrobial peptides in human pyelonephritis

**DOI:** 10.1101/2023.09.14.557715

**Authors:** Lars Borgards, Bente Siebels, Hannah Voss, Christoph Krisp, Jessica Schmitz, Lisa Schwarz, Devon Siemes, Philippa Spangenberg, Jan H Bräsen, Sibylle von Vietinghoff, Hartmut Schlüter, Florian Wagenlehner, Daniel R. Engel, Olga Shevchuk

## Abstract

Pyelonephritis (PN) is a frequent bacterial infection of the kidney and is often associated with severe diseases, organ loss and sepsis. Antibiotics are the cornerstone of therapy, however, increasing antibiotic resistance threatens therapy success and necessitates novel treatment strategies. Various proteins, such as antimicrobial peptides (AMPs), are key molecules of the innate immune response and insights into their regulation may help overcome multi-drug resistance and severe diseases. Using label-free liquid chromatography-tandem mass spectrometry (LC-MS/MS), several cellular, biological, and metabolic processes important for the antimicrobial response were identified, including a significant increase in previously undescribed proteins in human PN with antimicrobial function. Among others, we observed elevation of AMPs, such as calprotectin, azurocidin-1, and cathepsin G in the kidney, which we validated in the urine. Additionally, we observed a negative correlation of azurocidin-1 with plasma levels of C-reactive protein suggesting that the presence in the kidney may protect from severe diseases and systemic inflammation. This study represents the first renal proteomic dataset of human PN, enabling novel insights into the expression of AMPs in the context of PN.

**Lay Summary:** Growing antimicrobial resistance necessitates a better understanding of the expression of proteins that are critical for the immune response. Using mass spectrometry we identified AMPs in the kidney and urine of PN patients. Elevated levels of the AMP azurocidin-1 was associated with reduced systemic inflammation, indicated by lower C-reactive protein. Overall, this study identified expression of previously undescribed AMPs in the context of human PN. These proteins may play a pivotal role in protection from severe diseases and systemic inflammation.

## Introduction

Pyelonephritis (PN) is a frequent infection that results in severe inflammation of the renal parenchyma, causing severe morbidity, especially among elderly and immunodeficient patients. Potential long-term effects of PN include hypertension, proteinuria, renal fibrosis, and kidney failure [1, 2]. Gram-negative bacilli, such as *Escherichia coli* (40 – 80%) and *Klebsiella* spp. (10 – 25%), are the most frequent causative pathogens [1-4]. Infectious diseases of the urinary tract are commonly treated with antibiotics. However, multi-drug-resistant strains of uropathogenic bacteria become an increasingly urgent problem and necessitate the need for optimized treatment strategies and novel therapeutics [5]. For instance, 17 – 36% of patients with urosepsis are colonized with causative pathogens resistant against the initial empiric antibiotic treatment [6], requiring optimized treatment strategies.

As a first line of defense, the innate immune response plays a fundamental role in protecting the urothelium and renal parenchyma from pathogens [7]. Leukocytes and epithelial cells mediate innate immunity through direct phagocytosis of the pathogen or the release of soluble factors, such as components of the complement system, chemokines and cytokines, but also antimicrobial peptides and proteins (AMPs) [8] [9, 10]. AMPs are an integral part of the immune response within the urinary tract, as they exert a wide variety of microbicidal and immunoregulatory functions [8]. In addition to the direct inactivation of bacterial pathogens, e.g., by disrupting the bacterial cell membrane, AMPs modulate immunogenic activities, enhance chemokine production, regulate epithelial cell differentiation, and alter angiogenesis [8]. Thus, AMPs have attracted increased attention as prognostic markers and novel therapeutics [11].

Previous investigations of urinary tract infections (UTI) have mainly focused on AMPs in the urine of patients or animal models [12-14]. So far, multiple AMPs have been identified in the context of urinary tract immunity, including cathelicidin antimicrobial peptide (LL-37 or CAMP) [15], different α-and β-defensins [16], members of the ribonuclease A superfamily, such as RNase 7 [17], pentraxin-related protein 3 (PTX3) [18], uromodulin (UMOD) [19, 20], and metal-binding AMPs, such as lipocalin-2 [20-22], lactotransferrin (LTF) [14], and the S100A8/S100A9 heterodimer calprotectin [23-26]. The urothelium of the bladder and epithelial cells of the kidney and ureter were identified as primary sources for AMPs in the urinary tract, alongside cells of the innate immune system [10, 27]. Furthermore, within the kidney the α-intercalated cells of the collecting duct and thick ascending limb cells play a key role in the production and release of antimicrobial factors [10, 27]. However, data on the regulation of AMPs in the human kidney during PN is lacking. In this study, we generated a comprehensive LC-MS/MS-based dataset of the renal proteome in PN, providing a repository on the proteomic alterations in human PN. We observed elevation of certain AMPs in the kidney and validated the secretion of specific AMPs into the urine of human PN patients using antibody-based assays.

## Materials and methods

### Kidney samples of human PN patients

PN nephrectomy specimens (*n*=10; 20% men; mean age: 51.8 years; range: 22 – 76 years) and healthy sections from tumor nephrectomies (*n*=10; 30% men; mean age: 48.4 years; range: 7 – 82 years) were identified form the archives of the Nephropathology Unit at Hannover Medical School (MHH). Ethic MHH: No. 10183_BO-K_2022. The underlying diagnoses in patients undergoing nephrectomy are summarized in Supplementary Table 1.

### Urine and plasma samples of PN patients

Urine and plasma samples were collected from patients with acute PN (*n*=30) and healthy donors (*n*=10) by the Clinic for Urology, Pediatric Urology and Andrology of the Justus-Liebig-University Giessen. The ethical approval has been received by the independent ethics commission of the University Giessen, Germany (Ethical vote 280/20) on January 6^th^, 2021. The clinical data of pyelonephritis patients and control urine donors are summarized in Supplementary Table 3. The cases of acute pyelonephritis were clinically diagnosed, following the FDA’s definitions used for studies in the indication “Developing drugs for treatment. Guidance for industry, June 2018” (https://www.fda.gov/media/71313/download). Patients admitted to the urological department at the University Hospital Giessen with at least two of the following signs or symptoms were included: chills, rigors, or warmth associated with fever; flank pain; nausea or vomiting; dysuria; urinary frequency; urinary urgency; costo-vertebral angle tenderness upon physical examination; or a urine specimen with evidence of pyuria. Risk factors for recurrent UTI were categorized by urologists according to the ORENUC classification system [28]. The name ORENUC is an acronym standing for O (no known factors), R (risk for recurrent UTI), E (extra urogenital risk factors), N (nephropathy), U (urological risk factors that can be resolved by therapy), and C (catheter and the risk factors that cannot be resolved by therapy). For plasma preparation, 15 mL of blood were mixed with 15 mL PBS before adding the suspension to a 50 mL tube containing 15 mL Ficoll. The solution was centrifuged at 1,500 g for 10 min. Urine creatinine and C-reactive protein were measured in the central laboratory at the University Hospital Giessen. Following centrifugation, the plasma layer was transferred to cryotubes and stored at −20°C. Urine samples were centrifuged at 5,000 g for 10 min at 4°C and the supernatant was collected for ELISA of AMPs.

### Enzyme-linked immunosorbent assay (ELISA)

For analysis of the urinary levels of calprotectin, cathepsin G, azurocidin-1, and uromodulin by ELISA, 100 µL of sample or standard were transferred to a pre-coated 96-well-plate. The assay was performed according to the manufacturer’s instructions with a single technical replicate per sample (Uromodulin Human ELISA, BioVendor R&D, Cat. No.: RD191163200R; Azurocidin ELISA Kit, antibodies-online.com, Cat. No.: ABIN4881912; Cathepsin G ELISA Kit, antibodies-online.com, Cat. No.: ABIN6954455; IDK® Calprotectin (MRP 8/14) (Urine) ELISA, Immundiagnostik, Cat. No.: K 6928). Absorbance was measured on an iMark™ Microplate Absorbance Reader (Bio-Rad) at a wavelength of 450 nm. Calculated concentrations were normalized to creatinine values.

### Protein extraction and tryptic digestion for LC-MS/MS

60 µm FFPE tissue sections were diluted in 100 µL N-heptan and centrifuged for 5 min at 16,000 g. The supernatant was discarded, 100 µL ethanol were added and the samples were centrifuged again for 5 min at 16,000 g. Both steps were repeated once. Samples were diluted in 150 µL of 0.1 M tetraethylammonium bromide (TEAB) and 1% (w/v) sodium deoxycholate (SDC) buffer and heated to 95°C for 1h. Sonification was performed for six pulses at 30% power. The concentration of denatured proteins was determined by the Pierce™ BCA protein assay kit following the manufacturer’s instructions. 20 µg of protein for each sample were diluted in 50 µl SDC buffer. Disulfide bonds were reduced with 10 mM DTT at 60°C for 30 min. Cysteine residues were alkylated with 20 mM iodoacetamide (IAA) for 30 min at 37°C in the dark. Following tryptic digestion, the inhibition of trypsin activity as well as the precipitation of SDC were achieved by adding 1% formic acid (FA). Afterwards, samples were centrifuged for 5 min at 16,000 g. The supernatant was dried in a SpeedVac™ vacuum concentrator and the samples were stored at −20°C until further use. Prior to mass spectrometric analysis, peptides were resuspended in 0.1% FA to a final concentration of 1 µg/µl. 1 µg was used for LC-MS/MS acquisition.

### LC-MS/MS acquisition

LC-MS/MS measurements were performed on a quadrupole-ion-trap-orbitrap MS (Orbitrap Fusion, Thermo Fisher) coupled to a nano-UPLC (Dionex Ultimate 3000 UPLC system, Thermo Fisher). Chromatographic separation of peptides was achieved with a two-buffer system (buffer A: 0.1% FA in H_2_O, buffer B: 0.1% FA in ACN). Attached to the UPLC was a peptide trap (180 µm x 20 mm, 100 Å pore size, 5 µm particle size, Symmetry C18, Waters) for online desalting and purification, followed by a 25 cm C18 reversed-phase column (75 µm x 200 mm, 130 Å pore size, 1.7 µm particle size, peptide BEH C18, Waters). Peptides were separated using an 80 min gradient with linearly increasing ACN concentration from 2% to 30% ACN in 65 minutes. Eluting peptides were ionized using a nano-electrospray ionization source (nano-ESI) with a spray voltage of 1,800, transferred into the MS, and analyzed in data dependent acquisition (DDA) mode. For each MS1 scan, ions were accumulated for a maximum of 120 milliseconds or until a charge density of 2 x 10^5^ ions (AGC Target) was reached. Fourier-transformation based mass analysis of the data from the orbitrap mass analyzer was performed covering a mass range of 400 – 1,300 m/z with a resolution of 120,000 at m/z = 200. Peptides with charge states between 2+ −5+ above an intensity threshold of 1,000 were isolated within a 1.6 m/z isolation window in Top Speed mode for 3 seconds from each precursor scan and fragmented with a normalized collision energy of 30% using higher energy collisional dissociation (HCD). MS2 scanning was performed, using an ion trap mass analyzer at a rapid scan rate, covering a mass range of 380 – 1,500 m/z and accumulated for 60 ms or to an AGC target of 1 x 10^5^. Already fragmented peptides were excluded for 30 s.

### Raw MS data processing and statistics

LC-MS/MS data were searched with the Sequest algorithm integrated into the Proteome Discoverer software (v2.41.15, Thermo Fisher Scientific) against a reviewed human Swissprot database, obtained in April 2021, containing 20,365 entries. Carbamidomethylation was set as a fixed modification for cysteine residues. The oxidation of methionine, and pyro-glutamate formation at glutamine residues at the peptide N-terminus, as well as acetylation of the protein N-terminus were allowed as variable modifications. A maximum number of two missing tryptic cleavages was set. Peptides between six and 144 amino acids were considered. A strict cutoff (FDR < 0.01) was set for peptide and protein identification. Quantification was performed using the Minora Algorithm, implemented in Proteome Discoverer v2.4. Obtained protein abundances were log_2_-transformed and normalized through LOESS normalization [29].

For statistical analysis we considered 2,562 proteins, which were detected in at least one condition with at least two biological replicates. For each protein we calculated the fold change, *p*-value and signal-to-noise ratio. Subsequently, the *p*-value were adjusted for the FDR with Benjamini-Hochberg (*q*-value). Significant regulation was considered for proteins with log_2_(FC) >± 2 and a *q*-value < 0.05. Among 2,562 proteins 160 (6,2%) were upregulated and 507 (19,8%) downregulated. Cytoscape (Cytoscape_v3.8.0, ClueGO_v2.5.7; gene ontology databases from 08.05.2010; min GO level = 3; max GO level = 8; number of genes = 3; min percentage = 3.0; kappa score threshold = 0.4) was used for functional enrichment analysis.

The Pearson correlation Clusterheatmap was generated in python with plotly (v5.5.0) using 1273 proteins with a *q*-value < 0.05. Pearson correlation was calculated in pandas (v1.3.5) and the clustering was computed with scipy (v1.7.3). Regression plots were generated using the regplot method of the Seaborn (v.0.11.2) data visualization library for Python.

## Results

### LC-MS/MS reveals substantial alterations of the renal proteome in human PN

To resolve the global proteomic alterations caused by PN we compared FFPE tissue from PN nephrectomy specimens with healthy kidney tissue from tumor nephrectomies. HE-stained tissue sections indicated similar proportions of medullary and cortical tissue with more inflammation and less tubular integrity in PN (Figure 1 A-B, Supplementary Figure 1). To understand the underlying molecular determinants that modulate kidney inflammation and the anti-bacterial response during PN, we analyzed renal tissue sections with a DDA LC-MS/MS label-free proteomic quantitative approach. Collectively, more than three thousand proteins were identified with a false discovery rate (FDR) less than 1% (Supplementary Table 2). The highest correlation between protein expression was observed within the control group (Pearson correlation r = 0.85 −0.97) (Figure 1C). Low heterogeneity was present within PN tissue samples (r = 0.55 −0.95) and the lowest correlation (r = 0.28 −0.91) was found between the control and PN groups (Figure 1C). Dimensionality reduction by principal component analysis (PCA) and subsequent clustering revealed separation of the control and PN patients (Figure 1D). Based on PCA, Pearson correlation and histopathological scoring the patient PN10 was identified as an outlier and excluded from subsequent statistical analyses.

**Figure 1.**
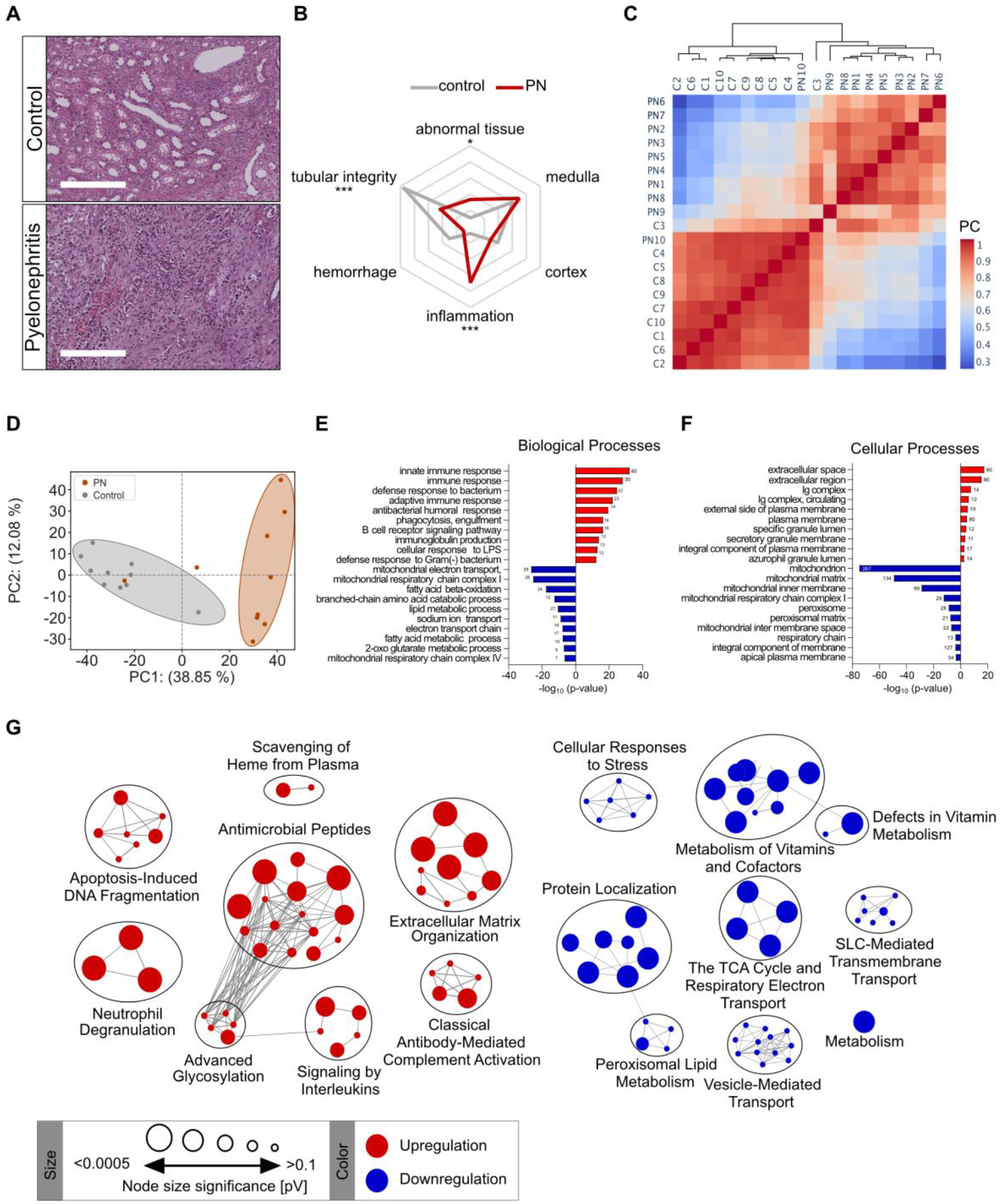
Proteomic alteration of the human kidney in PN. (**A, B**) Sections of PN (*n*=10) and control (C) kidneys (*n*=10) were stained with HE, scale represents 200 µm (A). Changes in the proportion of abnormal, medullary (medulla), and cortical (cortex) tissue, severity of inflammation (inflammation), hemorrhage and tubular integrity were evaluated using a scoring sheet with an arbitrary scale of 0-100. The average scoring of PN (red) and control (gray) samples are visualized using in a spider plot. *=*p*<0.05, \*\*\**p*<0.0005. (**C-G**) Label-free LC-MS/MS analysis of PN and control kidney sections. **(C)** Heatmap of Pearson correlations indicate similarity between donors. The Pearson correlation cluster heatmap was generated using 1,273 proteins that were detected in all 20 patients with a FDR-corrected *p*-value < 0.05. PC=Pearson correlation. (**D**) Principal component analysis (PCA) of the total kidney proteome of healthy and PN revealed condition-specific changes. (**E, F**) PN renal proteome reveals differential regulation of biological processes (E) and protein localization (F). Enrichment analysis by gene ontology of proteins expressed exclusively in one condition or with (log_2_(FC) > 2 or < −2, adjusted p-value < 0.05). (**G**) Distinct protein classes and pathways determined using the REACTOME database; Upregulated pathways are shown in red, downregulated pathways are shown in blue. The node size corresponds to the respective *p*-value.

Enrichment analysis using Gene Ontology Biological Process (GOBP) revealed upregulation of biological processes involved in the innate and adaptive immune response (Figure 1E). Among others, we observed upregulation of “*defense response to bacterium*” and “*antibacterial humoral response*” (Figure 1E), which also included “*AMP production*” (GO:0002775). At the same time, we observed a reduction of proteins involved in mitochondrial electron transport and assembly of the mitochondrial respiratory chain, fatty acid, and amino acid metabolism (Figure 1E). The Gene Ontology Cellular component (GOCC) indicated enrichment of proteins belonging to the *“extracellular space”, “plasma membrane”, “azurophil granules”*, and *“Ig complex”* in PN, whereas analysis of downregulated proteins revealed ontologies associated to mitochondria or peroxisomes (Figure 1F). Further classification of the proteomic changes by REACTOME-based overrepresentation analysis showed global upregulation of molecules involved in innate immunity, including factors of “*Neutrophil degranulation”*, “*Antimicrobial peptides”*, “*Extracellular matrix organization”*, “*Apoptosis-Induced DNA fragmentation”*, proteins involved in “*Signaling by Interleukins* as well as *Scavenging of Heme from Plasma”* (Figure 1G). We also observed down-regulation of metabolic pathways, such as “*Metabolism of Vitamins and Cofactors”* and *“TCA cycle and Respiratory Electron Chain*” (Figure 1G), Supplementary table 2). Thus, our data demonstrate global alterations of the proteome in human PN, particularly, upregulation of AMPs.

### AMPs constitute an enriched cluster in the PN kidney proteome

To gain insights into the kidney specific expression of AMPs in PN, we statistically analyzed the proteomic changes in the human PN and labeled the AMPs in a Volcano plot (Figure 2A). Among 16 AMPs detected in the kidney tissue, 13 were significantly regulated, such as S100A8, S100A9, cathepsin G (CTSG), neutrophil gelatinase-associated lipocalin (NGAL), lactotransferrin, eosinophil cationic protein (RNASE3), bactericidal permeability-increasing protein (BPI), azurocidin-1 (AZU1 or CAP37) and cathelicidin, as well as other proteins involved in the host defense response, e.g., neutrophil defensins, myeloperoxidase (MPO), and galectins. Resistin (RETN), which was recently identified as an antimicrobial peptide [30], was strongly induced in PN, whereas uromodulin was downregulated (Figure 2A).

**Figure 2.**
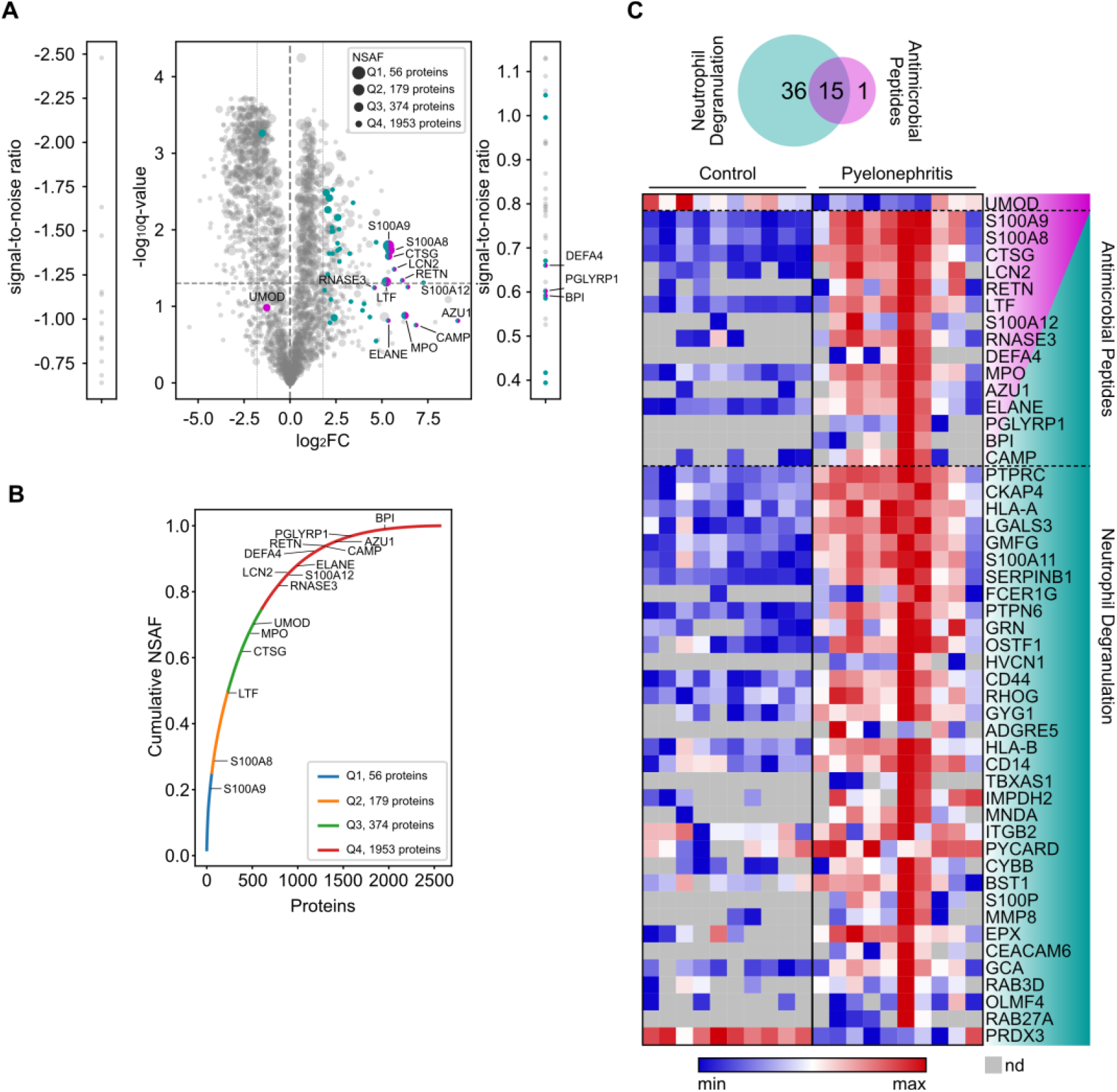
AMPs are overexpressed during PN. (**A**) Volcano plot depicting the log_2_ fold change (FC) and -log_10_ FDR-corrected *p*-value (*q*-value) of proteins measured by LC-MS/MS. Proteins detected only in one condition are indicated by signal-to-noise ratio. AMPs are highlighted in magenta; proteins involved in neutrophil degranulation are highlighted in cyan. Increasing abundance is indicated by the size of nodes using the normalized spectral abundance factor (NSAF) of the respective protein, quartile 1 (Q1) -Q4. (**B**) Cumulative NSAF values with specific annotation of AMPs. (**C**) Venn diagram and heat map of proteins curated to the REACTOME pathways *Neutrophil Degranulation* and *Antimicrobial Peptides*. The heatmap indicates the relative expression of these proteins; ordering according to signal-to-noise ratio and protein class was used. nd=not detected.

Additionally, we estimated the relative abundance of proteins in the renal tissue by calculating the normalized spectral abundance factor (NSAF). Based on NSAF values, some of the AMPs, e.g., S100A8 and S100A9 represented the bulk of the kidney proteome (Figure 2B). Most of these AMPs also belong to the class of neutrophil degranulation (Figure 2C), sequestering iron, or binding heme. These data represent a comprehensive dataset of AMPs in the kidney, identifying local expression of several AMPs that has not yet been described in the context of PN.

### Expression of AMPs in urine phenocopies the proteomic screening in the kidney

To study whether the AMPs from the proteomic screening are also expressed and regulated in the urine of PN patients, we collected urine from healthy (*n*=10) and PN donors (*n*=30), classified according to the ORENUC criteria, a stratification system for UTI [28]. Age, sex, estimated glomerular filtration rate (eGFR), and leukocyte counts in urine and plasma were determined (Table 1).

**Table 1.** *Characteristics and clinical data of urine donors. eGFR = estimated glomerular filtration rate, CRP-C-reactive protein, nd = not determined*.

| Parameters | PN | Control |
| --- | --- | --- |
| N | 30 | 10 |
| Mean age (min, max) | 38 (16 – 86) | 29 (23 – 31) |
| Sex (n) | ♀: 26; ♂: 4 | ♀: 6; ♂: 4 |
| eGFR | 101.6 (27.8 – 168.2) | nd |
| Fever [°C] | 39 (37.8 – 41.2) | nd |
| CRP [mg/mL] | 102.7 (13.7-280.2) | nd |
| Leukocytosis [cells/mL plasma] | 13.8 (8.9 – 25) | nd |
| Leukocyturia [cells/mL urine] | 2289.6 (31 – 33748) | nd |

To further evaluate the regulation of AMPs, S100A8 and S100A9, which comprise the heterodimer calprotectin and represented the most significantly regulated proteins, were analyzed by ELISA. Additionally, we evaluated the presence of azurocidin-1, uromodulin, and cathepsin G since data on the regulation of those AMPs in the human PN is lacking. Calprotectin, azurocidin-1 and cathepsin G were significantly upregulated in the urine of PN patients, whereas uromodulin remained unaltered (Figure 3A). Furthermore, we observed a positive correlation of calprotectin, azurocidin-1 and cathepsin G in urine (Figure 3B) and a negative correlation of azurocidin-1 with C-reactive protein (*p*-value=0.0274; Figure 3C). The urinary concentration of cathepsin G and calprotectin also showed a weak, negative correlation with plasma levels of CRP. However, correlation of cathepsin G and calprotectin with CRP, was not statistically significant with a *p*-value of 0.0709 and 0.0623, respectively. These data indicate that changes in the expression of AMPs in the kidney are reflected in urine and that increased azurocidin-1 expression is associated with decreased systemic inflammation.

**Figure 3.**
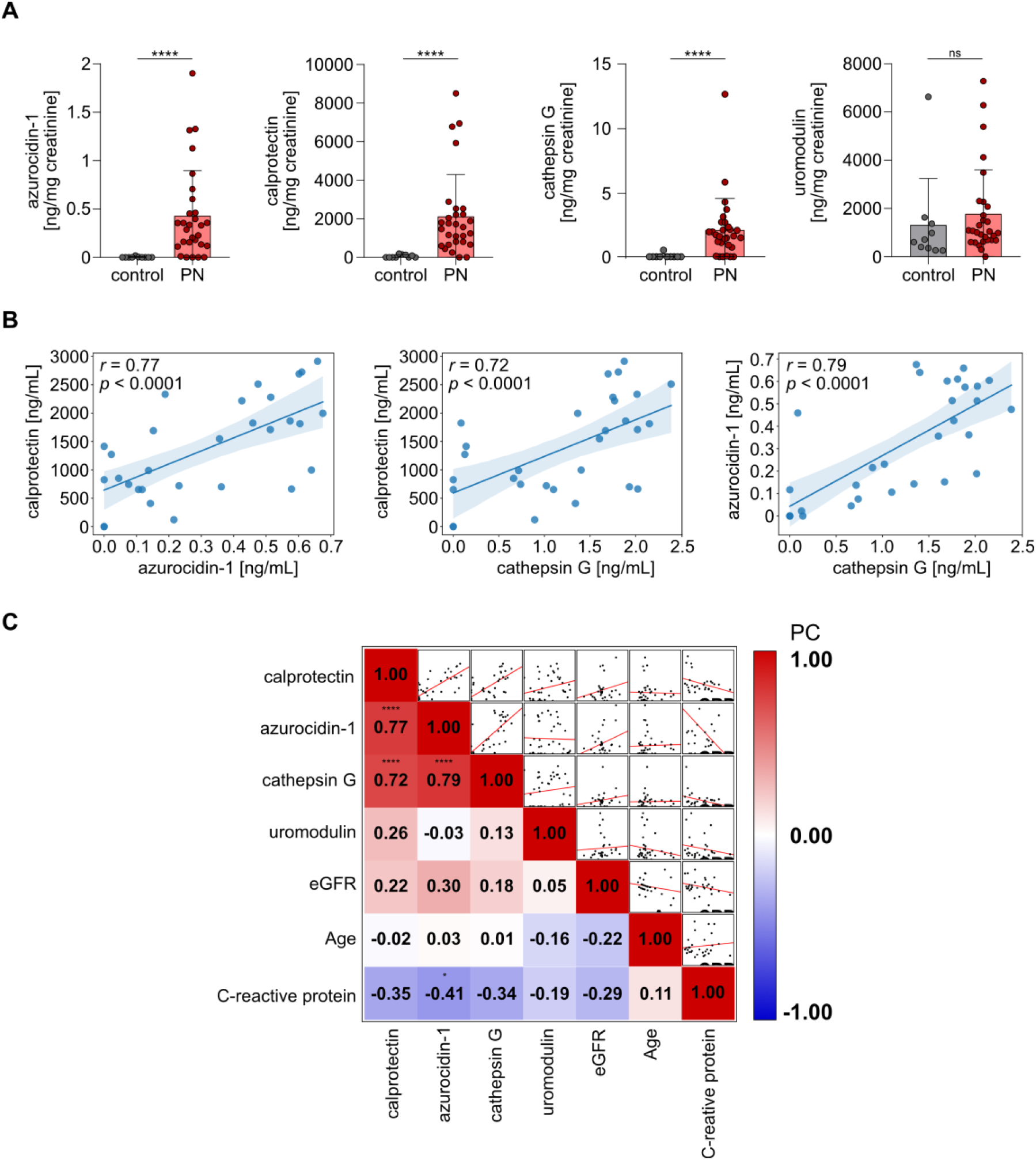
Upregulation of AMPs in the urine of acute PN patients. (**A**) Expression of calprotectin, azurocidin-1, cathepsin G, and uromodulin were measured by ELISA in urine from patients with acute PN (*n*=30) and healthy donors (*n*=10) and normalized to creatinine values. \*\*\*\**p*<0.0001, ns=not significant. (**B**) Regression plots show positive correlation of calprotectin, cathepsin G, and azurocidin expression levels, *r* indicates Pearson correlation, *p* indicates *p*-value. (C) Heat map of Pearson correlations (PC) between clinical parameters of urine donors. \**p*<0.05; \*\*\*\**p*< 0.0001.

## Discussion

PN is a severe bacterial disease associated with high morbidity and a growing prevalence of antibiotic resistance. To develop novel therapeutical approaches against PN, an understanding of the proteomic tissue microenvironment, and the expression of AMPs is required. In this study we found a pronounced adaptation of the renal proteome in PN, and substantial upregulation of several AMPs in the kidney and the urine. AMPs in the kidney were identified by LC-MS/MS and most of them could be annotated to the pathway “*Neutrophil Degranulation”* according to the Reactome database. The association of the detected AMPs with neutrophil degranulation suggests neutrophil recruitment, with these cells potentially serving as an important source of AMPs in PN. Despite the strong connection between the detected AMPs and neutrophils, other cell types have also been described to produce these molecules, such as epithelial cells in the case of cathelicidin [31], lactotransferrin [32], and lipocalin-2 [22] as well as monocytes and macrophages in the case of calprotectin [33, 34]. These aspects are currently not included in the curated databases used, i.e., REACTOME, and accordingly lead to the primary association to neutrophils.

We have found elevated levels of azurocidin-1, also known as cationic antimicrobial protein 37 (CAP37) or heparin-binding protein (HBP) in the kidney and urine of PN patients. Together with cathepsin G and neutrophil elastase, azurocidin-1 is a member of the serine protease family found in the specialized azurophilic granules of neutrophils [35, 36]. Crystallographic analysis indicates that the antibacterial activity of azurocidin-1 is likely mediated by a hydrophobic pocket (residues 20 to 44) that binds to the lipid A of Gram-negative bacteria [37]. During inflammation, this protein is released by neutrophils that arrive first at the site of infection and acts in a paracrine manner on endothelial cells, causing the formation of intercellular gaps and allowing leukocyte extravasation. Moreover, selective inactivation of azurocidin-1 prevented neutrophils from inducing endothelial hyperpermeability [38]. The lysosomal protease cathepsin G, which is also present in the azurophilic granules of neutrophils, was found to be strongly upregulated in PN. Previously, cathepsin G has been detected by LC-MS/MS in the urine pellet of UTI patients [39]. As a component of the neutrophil proteolytic machinery cathepsin G regulates the inflammatory response by stimulating the production of cytokines and chemokines [40]. Cathepsin G-specific hydrolysis of receptors, enzymes, cytokines, and other biologically active peptides leads to a modulation of chemotaxis and intercellular interactions, activation of the renin-angiotensin system (RAS), initiation of apoptosis, and many other processes [41]. Additional to proteolytic activity, cathepsin G possesses antimicrobial activity by encoding at least three antibacterial regions that by themselves can exert antibacterial action against Gram-negative and Gram-positive pathogens (peptides IIGGR and HPQYNQR; residues 1-5, 77-83 and 117-136 of full-length cathepsin G, respectively). Synthetic peptides corresponding to these sequences were found to exert broad-spectrum antimicrobial activity *in vitro*. They displayed conditions of temperature- and pH-dependent optima for antimicrobial action resembling that of the full-length enzyme [41, 42].

Iron acquisition systems are known to be essential for uropathogens, and UPEC strains encode highly diverse iron acquisition systems [43]. In line with this fact, iron depletion attenuates UPEC pathogenesis in a mouse and rat model of urinary tract infection and PN [44, 45]. We found elevated expression of several AMPs with iron sequestering or heme-binding properties, namely calprotectin, NGAL, lactotransferrin, myeloperoxidase and eosinophil peroxidase [46-49]. Calprotectin is known to be produced and released by neutrophils, monocytes, and macrophages [23] and was found to be upregulated in acute renal allograft failure, AKI [23], obstructive nephropathy (ON) [24], ischemia/reperfusion (I/R) injury [50], staphylococcus infection-associated glomerulonephritis (SAGN) [25], and anti-neutrophil cytoplasmic antibody (ANCA)-associated vasculitis (AAV) [26]. Despite its involvement in other kidney-related disorders, the effect of UTI on urinary and renal tissue concentrations of calprotectin remains elusive [51].

We observed that the expression of the AMPs did not correlate with the number of bacteria or leukocytes in the urine or blood but was negatively correlated with the plasma level of CRP, which is known to be elevated during the inflammatory response (Supplementary figure 2). Plasma CRP is produced by hepatocytes, predominantly under transcriptional control by the cytokine IL-6 [52], and a potential discriminative marker between cystitis and PN [53-55]. We speculate that higher levels of selective AMPs, i.e., azurocidin-1, are associated with a reduced systemic inflammatory response, whereas patients with reduced azurocidin-1 levels develop an IL-6-mediated acute-phase response, leading to CRP production.

An additional AMP of the urinary tract which has been identified in this study is uromodulin, also known as Tamm-Horsfall protein. Uromodulin is a GPI-anchored glycoprotein produced by epithelial cells of the thick ascending limb and distal tubule [27, 56]. Excreted via conserved proteolytic cleavage by hepsin [57, 58], uromodulin represents the most abundant protein in the urine. Data on uromodulin regulation suggest a protective role in urinary tract immunity. For instance, high urinary uromodulin levels have been shown to be associated with a lower risk of UTI in older adults [59]. Furthermore, patients who underwent kidney transplantation and recurrent UTIs had lower levels of uromodulin compared with persons without UTI, indicating an association between uromodulin levels and recurrence of infection. We observed slightly reduced uromodulin expression in the kidney during PN, though the alteration was not statistically significant. In opposite, uromodulin in the urine was slightly elevated in PN patients, which could indicate proteolytic excretion from the kidney. Its reduced expression in our dataset is in accordance with histological evaluations, which depict significantly reduced tubular integrity, as well as with a previous study describing uromodulin as a marker of kidney function and renal parenchymal integrity [60].

In addition to the aforementioned AMPs, we discovered strong upregulation of S100A12 (Calgranulin C) and Resistin. Calgranulin C has been shown to encode a 15-residue peptide at the C-terminus, named accordingly Calcitermin [61]. The peptide primarily targets Gram-negative bacteria and has optimal activity at acidic pH (pH 5.4), which represents the urine environment. Additionally, we observed strong expression of resistin, a cysteine-rich polypeptide hormone protein, which can directly kill bacteria by damaging their membranes [30]. Moreover, resistin can limit inflammation caused by microbial products, and modulate numerous host cellular responses to recruit and activate immune cells, promote the release of proinflammatory cytokines, reinforce interferon expression, and promote neutrophil extracellular trap formation [62, 63]. The expression of resistin is not only induced by microorganisms and pro-inflammatory cytokines but also regulated by vitamin A and its derivatives [64, 65], opening an avenue for dietary therapeutic regulation of this AMP in recurrent PN.

## Conclusion

This study provides the first renal proteomic dataset in human PN, yielding novel insights regarding the proteomic tissue adaptations in the context of this disease. A better understanding of the regulation of AMPs during PN may serve as the basis for designing novel, optimized treatment strategies for PN. Once this understanding is attained, it will pave the way for prospective translation studies. These efforts might be oriented towards the targeted activation of specific AMPs to address the issue of recurrent PN, or the treatment of patients utilizing short, bioactive peptides derived from the studied AMP proteins.

## Data availability

The raw mass spectrometric data and the PD result files have been deposited to the ProteomeXchange Consortium via the PRIDE partner repository with the dataset identifier: project accession: PXD042016 [66].

## Author Contributions

The clinical samples were collected and characterized by JS, FW, SVV, JHB. The experiments were performed by LB, OS. Mass spectrometry measurements, data analysis, visualization, and computational analysis were performed by BS, HV, CK, DS, PS, HS, OS. The study was written and proofread by LB, DRE and OS. All authors have read and commented on the article.

## Acknowledgement and funding

We acknowledge support by the Open Access Publication Fund of the University of Duisburg-Essen, the Central Animal Facilities of the Medical Faculty Essen, the Imaging Center Essen (Alexandra Brenzel and Dr. Anthony Squire) and the Immunoproteomics group (Dr. Olga Shevchuk, Stephanie Tautges-Schaefer, Stephanie Thiebes and Jenny Dick), supported by INST 20876/486-1 to DRE. DRE and OS received funding from the German Research Foundation: FOR5427 SP1 (OS); FOR5427 SP4 (DRE); EN984/15-1 and 16-1 (DRE); TR296 P09 and Z03 (DRE); TR332 A3 and Z1 (DRE).

## Supplementary material

**Supplementary Table 1:** Characteristics and clinical data of patients with partial kidney tissue resection or nephrectomy.

**Supplementary Table 2:** Normalized abundance of proteins identified by LC-MS/MS and corresponding statistics. Related to Figure 1C-G and Figure 2.

**Supplementary Table 3:** Clinical data of PN and control urine donors.

**Supplementary Figure 1:**
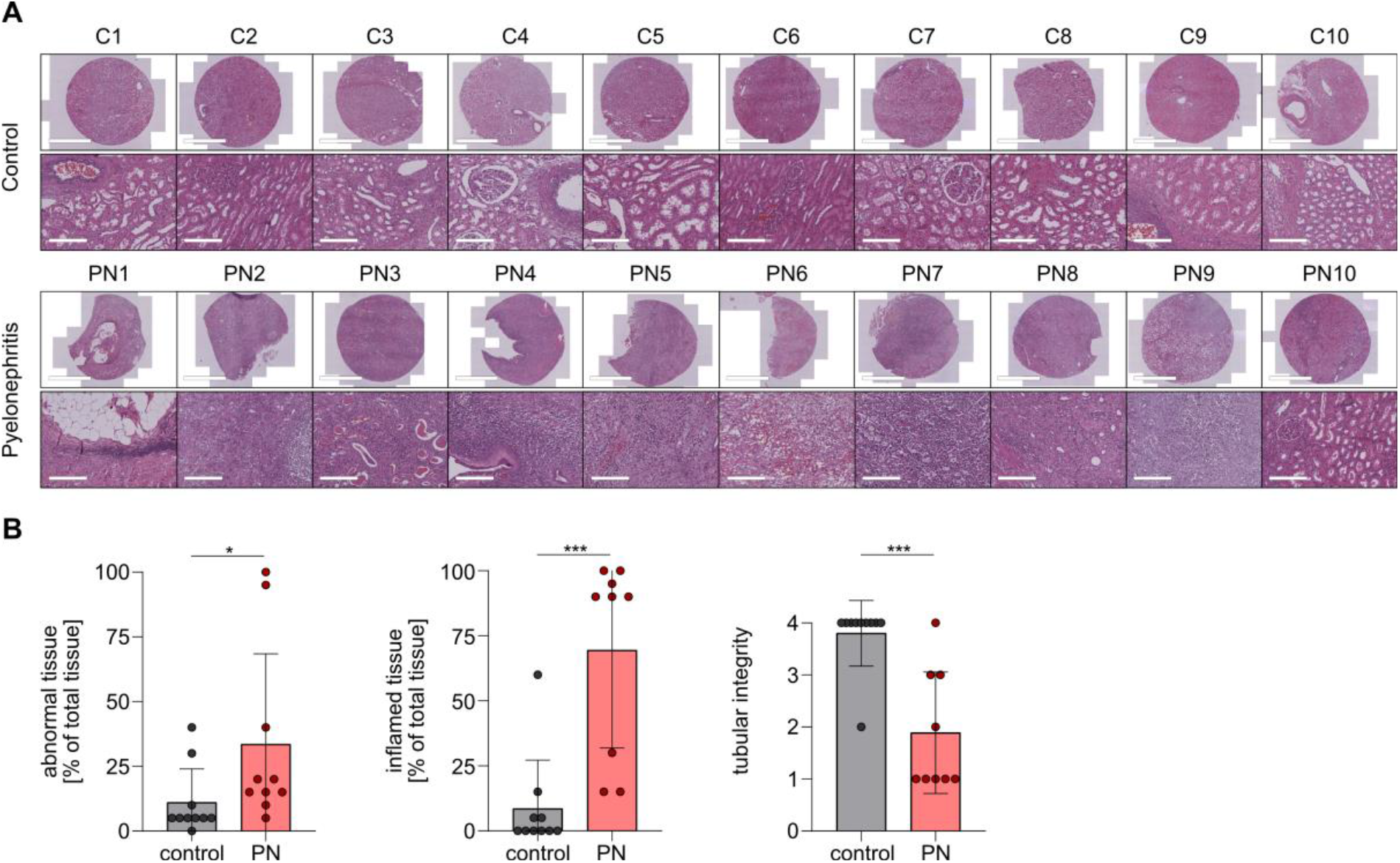
Evaluation of HE stained tissue from PN and control patients according to tissue abnormality, tissue inflammation and tubular integrity. Related to Figure 1 A, B. *=*p*<0.05, \*\*\**p*<0.0001.

**Supplementary Figure 2:**
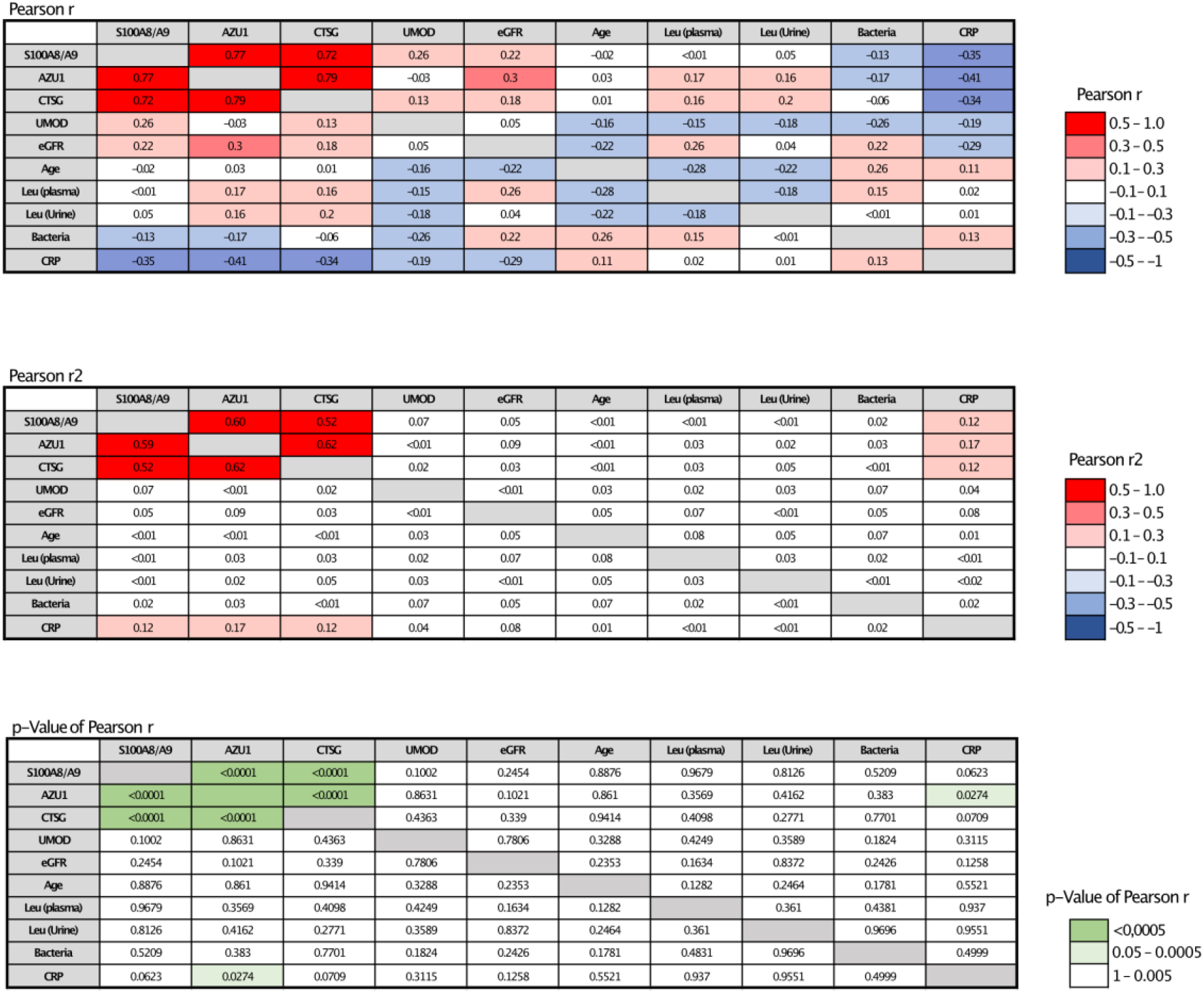
The Pearson correlation of AMPs and clinical data: Pearson r, r^2^, and p-value of each correlation.

## References

1. Tandogdu, Z. and F.M. Wagenlehner, Global epidemiology of urinary tract infections. Curr Opin Infect Dis, 2016. 29(1): p. 73–9.

2. Ferri, F.F., Ferri’s Clinical Advisor 2018 E-Book: 5 Books in 1. Ferri’s Clinical Advisor. 2017: Elsevier Health Sciences.

3. Stamm, W.E., et al., Urinary tract infections: from pathogenesis to treatment. J Infect Dis, 1989. 159(3): p. 400–6.

4. Talan, D.A., et al., Fluoroquinolone-Resistant and Extended-Spectrum beta-Lactamase-Producing Escherichia coli Infections in Patients with Pyelonephritis, United States(1). Emerg Infect Dis, 2016. 22(9): p. 1594–603.

5. Wagenlehner, F.M., et al., Non-Antibiotic Herbal Therapy (BNO 1045) versus Antibiotic Therapy (Fosfomycin Trometamol) for the Treatment of Acute Lower Uncomplicated Urinary Tract Infections in Women: A Double-Blind, Parallel-Group, Randomized, Multicentre, Non-Inferiority Phase III Trial. Urol Int, 2018. 101(3): p. 327–336.

6. Tandogdu, Z., et al., Antimicrobial resistance in urosepsis: outcomes from the multinational, multicenter global prevalence of infections in urology (GPIU) study 2003-2013. World J Urol, 2016. 34(8): p. 1193–200.

7. Song, J. and S.N. Abraham, Innate and adaptive immune responses in the urinary tract. Eur J Clin Invest, 2008. 38 Suppl 2: p. 21–8.

8. Becknell, B., et al., Amplifying renal immunity: the role of antimicrobial peptides in pyelonephritis. Nat Rev Nephrol, 2015. 11(11): p. 642–55.

9. Ching, C., et al., Innate immunity and urinary tract infection. Pediatr Nephrol, 2020. 35(7): p. 1183–1192.

10. Saxena, V., et al., Kidney intercalated cells are phagocytic and acidify internalized uropathogenic Escherichia coli. Nat Commun, 2021. 12(1): p. 2405.

11. Magana, M., et al., The value of antimicrobial peptides in the age of resistance. Lancet Infect Dis, 2020. 20(9): p. e216–e230.

12. Meylan, P.R., et al., Relationship between neutrophil-mediated oxidative injury during acute experimental pyelonephritis and chronic renal scarring. Infect Immun, 1989. 57(7): p. 2196–202.

13. Lertdumrongluk, K., et al., Diagnostic accuracy of urine heparin binding protein for pediatric acute pyelonephritis. Eur J Pediatr, 2015. 174(1): p. 43–8.

14. Oghumu, S., et al., Differential gene expression pattern in biopsies with renal allograft pyelonephritis and allograft rejection. Clin Transplant, 2016. 30(9): p. 1115–33.

15. Butler, D., et al., Immunomodulation therapy offers new molecular strategies to treat UTI. Nat Rev Urol, 2022. 19(7): p. 419–437.

16. Cox, S.N., et al., Formalin-fixed paraffin-embedded renal biopsy tissues: an underexploited biospecimen resource for gene expression profiling in IgA nephropathy. Sci Rep, 2020. 10(1): p. 15164.

17. Becknell, B., C. Ching, and J.D. Spencer, The Responses of the Ribonuclease A Superfamily to Urinary Tract Infection. Front Immunol, 2019. 10: p. 2786.

18. Jaillon, S., et al., The humoral pattern recognition molecule PTX3 is a key component of innate immunity against urinary tract infection. Immunity, 2014. 40(4): p. 621–32.

19. Reindl, J., et al., Uromodulin-related autosomal-dominant tubulointerstitial kidney disease-pathogenetic insights based on a case. Clin Kidney J, 2019. 12(2): p. 172–179.

20. Skowron, B., et al., Urinary neutrophil gelatinase-associated lipocalin, kidney injury molecule-1, uromodulin, and cystatin C concentrations in an experimental rat model of ascending acute kidney injury induced by pyelonephritis. J Physiol Pharmacol, 2018. 69(4).

21. Arambasic, J., et al., Differentiation of acute pyelonephritis from other febrile states in children using urinary neutrophil gelatinase-associated lipocalin (uNGAL). Clin Chem Lab Med, 2016. 54(1): p. 55–61.

22. Paragas, N., et al., alpha-Intercalated cells defend the urinary system from bacterial infection. J Clin Invest, 2014. 124(7): p. 2963–76.

23. Kim, A.J., et al., Klotho and S100A8/A9 as Discriminative Markers between Pre-Renal and Intrinsic Acute Kidney Injury. PLoS One, 2016. 11(1): p. e0147255.

24. Tammaro, A., et al., S100A8/A9 promotes parenchymal damage and renal fibrosis in obstructive nephropathy. Clin Exp Immunol, 2018. 193(3): p. 361–375.

25. Satoskar, A.A., et al., Differentiating Staphylococcus infection-associated glomerulonephritis and primary IgA nephropathy: a mass spectrometry-based exploratory study. Sci Rep, 2020. 10(1): p. 17179.

26. Pepper, R.J., et al., Leukocyte and serum S100A8/S100A9 expression reflects disease activity in ANCA-associated vasculitis and glomerulonephritis. Kidney Int, 2013. 83(6): p. 1150–8.

27. Tokonami, N., et al., Uromodulin is expressed in the distal convoluted tubule, where it is critical for regulation of the sodium chloride cotransporter NCC. Kidney Int, 2018. 94(4): p. 701–715.

28. Johansen, T.E., et al., Critical review of current definitions of urinary tract infections and proposal of an EAU/ESIU classification system. Int J Antimicrob Agents, 2011. 38 Suppl: p. 64–70.

29. Ballman, K.V., et al., Faster cyclic loess: normalizing RNA arrays via linear models. Bioinformatics, 2004. 20(16): p. 2778–86.

30. Li, Y., et al., Resistin, a Novel Host Defense Peptide of Innate Immunity. Front Immunol, 2021. 12: p. 699807.

31. Chromek, M., et al., What Keeps the Urinary Tract Sterile?: The Antimicrobial Peptide Cathelicidin Protects the Urinary Tract against Invasive Bacterial Infection. Nat Med 12: 636-640, 2006. J Am Soc Nephrol, 2006. 17(12): p. 3267–3272.

32. Abrink, M., et al., Expression of lactoferrin in the kidney: implications for innate immunity and iron metabolism. Kidney Int, 2000. 57(5): p. 2004–10.

33. Edgeworth, J., et al., Identification of p8,14 as a highly abundant heterodimeric calcium binding protein complex of myeloid cells. J Biol Chem, 1991. 266(12): p. 7706–13.

34. Odink, K., et al., Two calcium-binding proteins in infiltrate macrophages of rheumatoid arthritis. Nature, 1987. 330(6143): p. 80–2.

35. Morgan, J.G., et al., Cloning of the cDNA for the serine protease homolog CAP37/azurocidin, a microbicidal and chemotactic protein from human granulocytes. J Immunol, 1991. 147(9): p. 3210–4.

36. Zimmer, M., et al., Three human elastase-like genes coordinately expressed in the myelomonocyte lineage are organized as a single genetic locus on 19pter. Proc Natl Acad Sci U S A, 1992. 89(17): p. 8215–9.

37. Kastrup, J.S., et al., Two mutants of human heparin binding protein (CAP37): toward the understanding of the nature of lipid A/LPS and BPTI binding. Proteins, 2001. 42(4): p. 442–51.

38. Gautam, N., et al., Heparin-binding protein (HBP/CAP37): a missing link in neutrophil-evoked alteration of vascular permeability. Nat Med, 2001. 7(10): p. 1123–7.

39. Yu, Y., et al., Characterization of Early-Phase Neutrophil Extracellular Traps in Urinary Tract Infections. PLoS Pathog, 2017. 13(1): p. e1006151.

40. Gao, S., et al., Cathepsin G and Its Role in Inflammation and Autoimmune Diseases. Arch Rheumatol, 2018. 33(4): p. 498–504.

41. Meyer-Hoffert, U., Neutrophil-derived serine proteases modulate innate immune responses. Front Biosci (Landmark Ed), 2009. 14(9): p. 3409–18.

42. Bangalore, N., et al., Identification of the primary antimicrobial domains in human neutrophil cathepsin G. J Biol Chem, 1990. 265(23): p. 13584–8.

43. Frick-Cheng, A.E., et al., Ferric Citrate Uptake Is a Virulence Factor in Uropathogenic Escherichia coli. mBio, 2022. 13(3): p. e0103522.

44. Bauckman, K.A., et al., Dietary restriction of iron availability attenuates UPEC pathogenesis in a mouse model of urinary tract infection. Am J Physiol Renal Physiol, 2019. 316(5): p. F814–F822.

45. Guze, L.B., et al., Effect of iron on acute pyelonephritis and later chronic changes. Kidney Int, 1982. 21(6): p. 808–12.

46. Nakashige, T.G., et al., Human calprotectin is an iron-sequestering host-defense protein. Nat Chem Biol, 2015. 11(10): p. 765–71.

47. Goetz, D.H., et al., The neutrophil lipocalin NGAL is a bacteriostatic agent that interferes with siderophore-mediated iron acquisition. Mol Cell, 2002. 10(5): p. 1033–43.

48. Jacquet, A., et al., Site-directed mutants of human myeloperoxidase. A topological approach to the heme-binding site. FEBS Lett, 1992. 302(2): p. 189–91.

49. Wever, R., et al., Characterization of the peroxidase in human eosinophils. Eur J Biochem, 1980. 108(2): p. 491–5.

50. Dessing, M.C., et al., The calcium-binding protein complex S100A8/A9 has a crucial role in controlling macrophage-mediated renal repair following ischemia/reperfusion. Kidney Int, 2015. 87(1): p. 85–94.

51. van Zoelen, M.A., et al., Expression and role of myeloid-related protein-14 in clinical and experimental sepsis. Am J Respir Crit Care Med, 2009. 180(11): p. 1098–106.

52. Pepys, M.B. and G.M. Hirschfield, C-reactive protein: a critical update. J Clin Invest, 2003. 111(12): p. 1805–12.

53. Mazaheri, M., Serum Interleukin-6 and Interleukin-8 are Sensitive Markers for Early Detection of Pyelonephritis and Its Prevention to Progression to Chronic Kidney Disease. Int J Prev Med, 2021. 12: p. 2.

54. Al Rushood, M., A. Al-Eisa, and R. Al-Attiyah, Serum and Urine Interleukin-6 and Interleukin-8 Levels Do Not Differentiate Acute Pyelonephritis from Lower Urinary Tract Infections in Children. J Inflamm Res, 2020. 13: p. 789–797.

55. Mahyar, A., et al., Serum levels of interleukin-6 and interleukin-8 as diagnostic markers of acute pyelonephritis in children. Korean J Pediatr, 2013. 56(5): p. 218–23.

56. Lhotta, K., Uromodulin and chronic kidney disease. Kidney Blood Press Res, 2010. 33(5): p. 393–8.

57. Santambrogio, S., et al., Urinary uromodulin carries an intact ZP domain generated by a conserved C-terminal proteolytic cleavage. Biochem Biophys Res Commun, 2008. 370(3): p. 410–3.

58. Brunati, M., et al., The serine protease hepsin mediates urinary secretion and polymerisation of Zona Pellucida domain protein uromodulin. Elife, 2015. 4: p. e08887.

59. Garimella, P.S., et al., Urinary Uromodulin and Risk of Urinary Tract Infections: The Cardiovascular Health Study. Am J Kidney Dis, 2017. 69(6): p. 744–751.

60. Scherberich, J.E., et al., Serum uromodulin-a marker of kidney function and renal parenchymal integrity. Nephrol Dial Transplant, 2018. 33(2): p. 284–295.

61. Cole, A.M., et al., Calcitermin, a novel antimicrobial peptide isolated from human airway secretions. FEBS Lett, 2001. 504(1-2): p. 5–10.

62. Jiang, S., et al., Human resistin promotes neutrophil proinflammatory activation and neutrophil extracellular trap formation and increases severity of acute lung injury. J Immunol, 2014. 192(10): p. 4795–803.

63. Jang, J.C., et al., Human resistin protects against endotoxic shock by blocking LPS-TLR4 interaction. Proc Natl Acad Sci U S A, 2017. 114(48): p. E10399–E10408.

64. Propheter, D.C., et al., Resistin-like molecule beta is a bactericidal protein that promotes spatial segregation of the microbiota and the colonic epithelium. Proc Natl Acad Sci U S A, 2017. 114(42): p. 11027–11033.

65. Harris, T.A., et al., Resistin-like Molecule alpha Provides Vitamin-A-Dependent Antimicrobial Protection in the Skin. Cell Host Microbe, 2019. 25(6): p. 777–788 e8.

66. Perez-Riverol, Y., et al., The PRIDE database resources in 2022: a hub for mass spectrometrybased proteomics evidences. Nucleic Acids Res, 2022. 50(D1): p. D543–D552.

